# Ingredients in Victoria’s Secret Bombshell and Ivanka Trump eaux de parfums that repel mosquitoes

**DOI:** 10.1101/172304

**Authors:** Fangfang Zeng, Pingxi Xu, Kaiming Tan, Paulo H. G. Zarbin, Walter S. Leal

**Affiliations:** Department of Molecular and Cellular Biology, University of California-Davis, Davis, CA 95616, USA; Departamento de Química, Universidade Federal do Paraná, Laboratório de Ecologia Química e Síntese Orgânica, CP 19081 CEP 81531-990 Curitiba-PR, Brazil

**Keywords:** DEET, surface landing and feeding assay, methyl dihydrojasmonate, lilial, galaxolide, isopropyl myristate, lyral, CquiOR136, Bombshell, Ivanka Trump eau de parfum, southern house mosquito, *Culex quinquefasciatus*

## Abstract

Insect repellents are widely used to fend off nuisance mosquitoes and, more importantly, to reduce or eliminate mosquito bites in areas where viruses and other vector-borne diseases are circulating. Synthesized more than six decades ago, DEET is the most widely used insect repellent. Plant-derived compounds are used in a plethora of commercial formulations and natural recipes to repel mosquitoes. They are also used as fragrances. We analysed Bombshell^®^ to identify the constituent(s) eliciting a previously reported “off-label” repellence activity. The two major fragrance ingredients in Bombshell, i.e., methyl dihydrojasmonate and lilial, demonstrated strong repellence against the southern house mosquito, *Culex quinquefasciatus*, in laboratory assays. Both compounds activated a previously identified DEET-sensitive odorant receptor, CquiOR136. These compounds were also major constituents of Ivanka Trump eau de parfum. The methyl dihydrojasmonate content was higher in the Ivanka Trump perfume than in Bombshell, the reverse being true for lilial. Both Bombshell and Ivanka Trump eaux de parfums retained activity for as long as 6 hours in laboratory assays. Although wearing these perfumes may repel nuisance mosquitoes, their use as “off-label” repellents against infected mosquitoes is not recommended. A panel of 104 students (18-23 years old) conducted a blind test to compare the two eaux de parfums and showed a preference for Bombshell over Ivanka Trump’s brand, particularly among women.

## Introduction

Insect repellents are used not only as prophylactic tools for travellers to and people living in endemic or outbreak areas of malaria, dengue, chikungunya, Zika, West Nile fever, encephalitis, and other vector-borne diseases, but also for reducing bites by nuisance mosquitoes. A plethora of repellents are derived from plants (botanical repellents) and other natural sources [1-3], but the synthetic compound *N*,*N*-diethyl-3-methylbenzamide (DEET) is the most widely used insect repellent. In the United States, there are currently approximately 120 repellent formulations registered with the EPA for direct application on human skin that contain 4-99% DEET [4]. Significant parts of the population that can afford and should wear repellents do not use DEET, because of undesirable properties, such as unpleasant odor and reactivity with eyeglass frames and watchbands. Additionally, a group of natural product aficionados embrace the misleading notion that natural is safe and synthetic is harmful, so they too do not use DEET. As the old repellent on the market, DEET has been scrutinized more than any of its counterparts and has a remarkable safety record [5], but one has to consider that no chemicals are “absolutely safe.” One of the more modern alternatives to DEET is picaridin [5], which dermatologists recommend as a second-line agent after concluding, based on peer-reviewed literature, that DEET demonstrates a strong and consistent ability to reduce mosquito bites relative to other repellents [6]. In summary, DEET is considered safe, or strictly speaking, a low-risk, high-benefit repellent. However, its continuous application at high doses is a matter of concern given DEET’s high levels of skin penetration [7]. Therefore, the use of repellents mild on the skin, albeit less effective (e.g., citronella oil and other plant-derived compounds), may be an alternative for those attempting to reduce bites of nuisance mosquitoes, but a high-risk strategy for those needing protection against infected mosquitoes.

The fragrance industry too still uses plant materials as ingredients [8]. Recently, it was reported that a commercial perfume, Victoria’s Secret Bombshell^®^ eau de parfum (hereafter Bombshell), showed strong repellence against mosquitoes [9]. We then asked the questions what constituents (fragrances) in Bombshell^®^ contribute to its repellence effect and compared this perfume with another equivalent product in the market, specifically Ivanka Trump eau de parfum. Here, we report that the active ingredient in Bombshell responsible for repellence activity are a plant-derived compound and commonly used synthetic fragrance, methyl dihydrojasmonate (=Hedione^®^) and a synthetic aromatic aldehyde, commonly used in cosmetics, lilial. These fragrances are also major constituents of Ivanka Trump eau de parfum, and they both activate a mosquito odorant receptor sensitive to DEET, CquiOR136. In laboratory assays, the two eaux de parfums showed repellence activity comparable to that elicited by DEET for as long as 6 hours. Lastly, we conducted a blind test with young students and recorded a slight preference for Bombshell^®^ eau de parfum over Ivanka Trump eau de parfum by men, whereas women showed a more pronounced preference for Bombshell^®^.

## Materials and Methods

### Mosquitoes

The laboratory colony of *Cx. quinquefasciatus* used in this study (“Davis colony’) originated from mosquitoes collected in Merced, California in the 1950s. The original “Merced colony” has been maintained in the Kearney Agricultural Center (KAC), University of California by Dr. Anthon Cornel. The “Davis colony” was initiated from eggs of the “Merced colony” provided by Dr. Anthon Cornel and has been maintained at Davis for more than six years under a photoperiod of 12:12 h (L:D), 27±1°C, and 75% relative humidity.

### Chemicals

DEET (PESTANAL^®^ analytical standard grade, 99.5%) was acquired from Sigma-Aldrich (catalogue number, 36452-250MG). Methyl dihydrojasmonate (>98%) and galaxolide (50% in isopropyl myristate) were from Bedoukian Research Inc. (Danbury, CT, USA). Lilial (=Lysmeral® EXTRA, code #503750) was acquired from Vigon International (East Stroudbsburg, PA, USA). Other chemicals, including isopropyl myristate (catalogue #172472, 98%) lyral (=4-(4-hydroxy-4-methyl)-3-cyclohexene-1-carboxaldehyde, catalogue #95594, >97%), galaxolide (analytical standard, >85%), were acquired from Sigma-Aldrich (Milwaukee, WI, USA). Victoria’s Secret Bombshell eau de parfum and Ivanka Trump eau de parfum spray vaporisateur were acquired from Amazon.com.

### Chemical analyses

Gas chromatography-mass spectrometry (GC-MS) analyses were performed on a 5973 Network Mass Selective Detector linked to a 6890 Series GC System Plus+ (Agilent Technologies, Palo Alto, CA), which was equipped with an HP-5MS capillary column (30 m x 0.25 mm; 0.25 μm film; Agilent Technologies). The oven temperature was set at the initial temperature of 70°C, for 1 min then the temperature was raised at a rate of 10°C/min to 270°C, and held at this final temperature for 10 min. After each run, the oven temperature was held at 290°C for 10 min; in short, 70°C (1)-10°C/min-270 (10); post run, 290°C (10). The injector was operated at 250°C in a pulsed splitless mode (18.5 psi for 1.5 min; purge flow, 50 ml/min, 1.5 min; saver flow, 20 ml/min, 2 min). MS transfer line was set at 280°C, MS quad and MS sources were set at 150°C and 230°C, respectively. GC coupled with Fourier transform infrared spectroscopy (GC-FT/IR) was carried out on a Shimadzu GC2010, coupled to a DiscovIR-GC infrared detector (DANI Instruments, Marlborough, Massachusetts, USA), with a scan range of 4000-750 cm^-1^ and resolution of 8 cm^-1^. The GC was equipped with an RTX-5 capillary column (30 m × 0.25 mm × 0.25 µm film thickness; Restek, Bellefonte, PA, USA), and injections were performed in splitless mode at 250°C (injector temperature). The column temperature was programmed to start at 50°C for 1 min and subsequently increased to 250°C at a rate of 7°C min^-1^ with a final hold of 10 min. Quantification was done on a gas chromatograph 6890 Series GC (Agilent Technologies), equipped with an HP-5MS column (same dimensions), with the following program for the oven temperature 70°C (1)-10°C/min-290 (5); post-run 290°C (5). The injector was operated at 250°C and in pulsed splitless mode (30 psi for 1 min; purge flow 41.7 ml/min for 1 min, and gas saver at 20 ml/min, 3 min). The response of the flame ionization detector (FID), which operated at 250°C, was calibrated by injecting multiple times (n≥3) standard samples of methyl dihydrojasmonate and lilial and measuring the areas of the peaks. Linear regression analysis from the data generated with injections of 25, 50, 100, and 200 ng of methyl dihydrojasmonate gave the equation Y (amount in ng) = 0.072X-1.518 (R^2^ = 0.986; F = 138.4; P = 0.007); X = measured area. Likewise, linear regression analysis of peak areas vis-à-vis injections of 10, 25, 50, and 100 ng of lilial generated the following equation: Y (amount in ng) = 0.053 X + 3.085 (R2 = 0.999; F = 2298; P = 0.0004). These equations were used to estimate the contents of methyl dihydrojasmonate and lilial in samples of Bombshell and Ivanka Trump eaux de parfums (n=3 each).

### Sample preparations and other procedures

For GC-MS analyses, samples were prepared in hexane and dried up with anhydrous sodium sulphate to eliminate traces of water derived from the perfumes. For GC analysis/quantification, samples were prepared in ethanol. Stock solutions (10%) were diluted in decadic steps from 10% to 0.1%_m/v_. One microliter of 0.1% solutions were injected to estimate the concentrations of methyl dihydrojasmonate and lilial dispensed from the perfume vials. To estimate the density of these perfumes, we weighted in triplicate the amount of each perfume in 25 μl capillary tubes (Drummond Scientific Company, Broomall, PA, USA). After placing one capillary inside a 4-ml glass on an analytical balance scale (GA 110 Electronic Laboratory Balance Scale, Ohaus Corporation, Parsippany, NJ, USA), the balance was zeroed, the capillary tube was filled with the test perfume, and the amount weighted. To estimate the amount of perfume dispensed per spray and the area of the body covered, a bottle of each perfume was held at about 10 cm from the forearm and the area covered by a single spray was measured. Then, the same procedure was done at a short distance to collect the entire spray into a 4-ml glass vial, which was weighted in an analytical balance.

### Behavior measurement

An improved version of the surface landing and feeding behavioral assay has been described in detail elsewhere [10]. In short, a two-choice arena was constructed in which two Dudley tubes painted inside with black ink protrude inside of a mosquito cage. With water at 28°C circulating inside these tubes, their ends serve not only as physical stimuli (colour and temperature), but also to hold dental cotton rolls. Syringe needles on the top of these tubes delivered carbon dioxide (at 50 ml/min) and held cotton rolls in place. Insect pins placed 1.8 cm above the syringe needles held filter paper rings (width 4 cm; 25 cm; overlapped 1 cm for stapling), which served as a spatial repellent source (and control). Defibrinated sheep blood (100 μl) was loaded on dental cotton rolls and one was placed on each side of the arena. Each filter paper was loaded with 200 μl of test sample or solvent and placed in the treatment or control arena, respectively, and tested soon after solvent evaporation (1-2 min). For the protection time experiments, samples and control were prepared and tested soon after solvent evaporation (t = 0 h), 2, 4, and 6 h after the sample preparations (t = 2, 4, and 6 h, respectively). In these cases, samples were prepared in advance to start all experiments at the beginning of the scotophase with aged samples. Responses of sugar-fed, blood-seeking, 5- to 7-day-old female mosquitoes were recorded for 5 min with a Super NightShot Plus infrared camcorder (Sony Digital Handycam, DCR-DVD 810). The number of mosquitoes that landed and continued to feed on each side of the arena was recorded as an endpoint measurement. Females on treatment and control sides of the arena were gently removed with a high finish pointed brush, and treatment and control sides were inverted before a new trial was initiated.

### Statistical analysis

Data from the surface landing and feeding assay were transformed (arcsin of response fractions) before paired two-tailed Student *t* test comparisons. For clarity, data are expressed as mean ± SEM. Data related to repellence over time are expressed in terms of protection rate, following WHO and EPA recommendations. Thus, P % = [1 –(T/C)] x 100, where C and T are the number of mosquitoes responding to the control and treated (repellent) side of the arena. In both cases, percentages were calculated with Excel spread sheets for subsequent analyses with Prism7 (GraphPad, La Jolla, CA). Data that did not meet the assumption of normality (Shapiro-Wilk test) were analyzed using the Mann-Whitney, two-tailed test.

### Two-electrode voltage clamp records

The two-electrode voltage-clamp (TEVC) technique was used to measure odorant-induced currents in the *Xenopus* oocyte recording system, with a holding potential of -80 mV. Signals were amplified with an OC-725C amplifier (Warner Instruments, Hamden, CT, USA), low-pass-filter at 50 Hz, and digitized at 1 kHz. Data acquisition and analyses were conducted with Digidata 1440A and pCLAMP software (Molecular Devices, Sunnyvale, CA, USA). Responses of CquiOR136/CquiOrco-expressing oocytes to DEET, methyl dihydrojasmonate, and lilial were compared at the same dose (1 mM, n = 5) and using different oocytes (n = 3).

### Fragrance preferences

A blind test was conducted with students (18-23 years old) leaving biochemistry classes on the UC-Davis campus in the winter quarter of 2017. Students were asked if they would volunteer to compare two fragrances, which were presented in spray bottles (Clear Boston Ground Bottle with Atomizer, BRF1AB, specialtybottle.com) labelled with the following code names: Isoleucine/Threonine (IT for Ivanka Trump eau de parfum) and Serine/Histidine (SH, for Bombshell). Students were asked if they preferred one of these two perfumes and were provided with an optional column to make “other remarks” and disclose the tester’s gender. To optimize the number of participants between classes, aliquots of the two eaux de parfums were transferred to three bottles each.

## Results and Discussion

Gas chromatography-mass spectrometry analyses showed that the top four major constituents of Bombshell^®^ were methyl dihydrojasmonate, lilial, galaxolide, and isopropryl myristate (Fig. 1). The diastereomers (=diastereoisomers) of methyl dihydrojasmonate [IUPAC name: methyl 2-(3-oxo-2-pentylcyclopentyl)acetate] appeared at 13.34 and 13.62 min. Not surprisingly, their mass spectral data (base peak, m/z 83; M^+^ = 226; other significant fragment, m/z 153) and GC-FT/IR data (C=O stretching, 1737 cm^-1^, strong C-H stretching, 2957 cm^-1^, weak) were indistinguishable from those obtained with an authentic sample of methyl dihydrojasmonate. Although methyl dihydrojasmonate is a natural product, occurring in trace amounts in tea flavour, Lima orange, and apparently in several other fruits and flowers [11], it is a well-known synthetic fragrance, also called Hedione^®^, which was first prepared in the early 1960s for partial confirmation of the proposed structure of methyl jasmonate [11]. The peaks of both authentic lilial and the fragrance from Bombshell (Fig. 1) appeared at the same retention time (11.84 min) and their mass spectra (MS) were identical (base peak, m/z 189; M^+^ = 204). GC-FT/IR data showed the characteristic bands at 1722 (strong) and 2966 cm^-1^ (strong) [12]. Likewise, we identified isopropyl myristate and galaxolide by comparison with authentic samples.

**Fig. 1.**
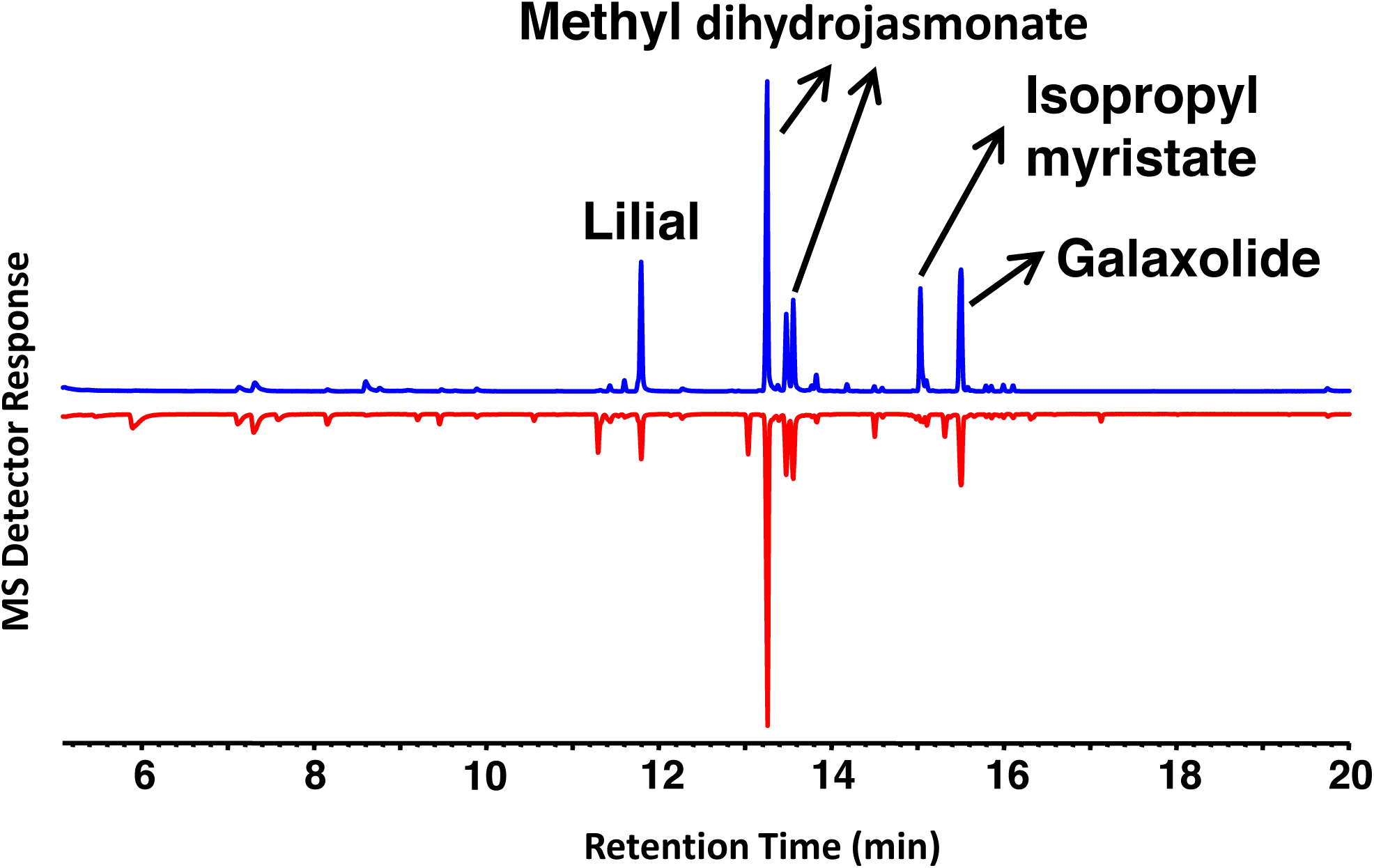
MS chromatogram profiles of Bombshell^®^ (upper trace in blue) and Ivanka Trump (lower trace in red) eaux de parfums. The peaks for the four major constituents of Bombshell are labelled. Three of them are the major constituents in the Ivanka Trump eau de parfum.

Next, we tested whether these individual compounds were repellents in our surface landing and feeding assay [13]. For this, we used DEET at 1% as a positive control and the southern house mosquito, *Culex quinquefasciatus*, as test mosquitoes. Both methyl dihydrojasmonate (MDJ) (Fig. 2A) and lilial at 1% (Fig. 2B) showed strong repellence activity. By contrast, neither isopropyl myristate (IM) (Fig. 2C) nor galaxolide (Fig. 2D) repelled *Culex* mosquitoes. When tested in 2:1 mixtures at 1 and 5%, methyl dihydrojasmonate and lilial did not have a synergistic effect (Fig. 3A and B, respectively).

**Fig. 2.**
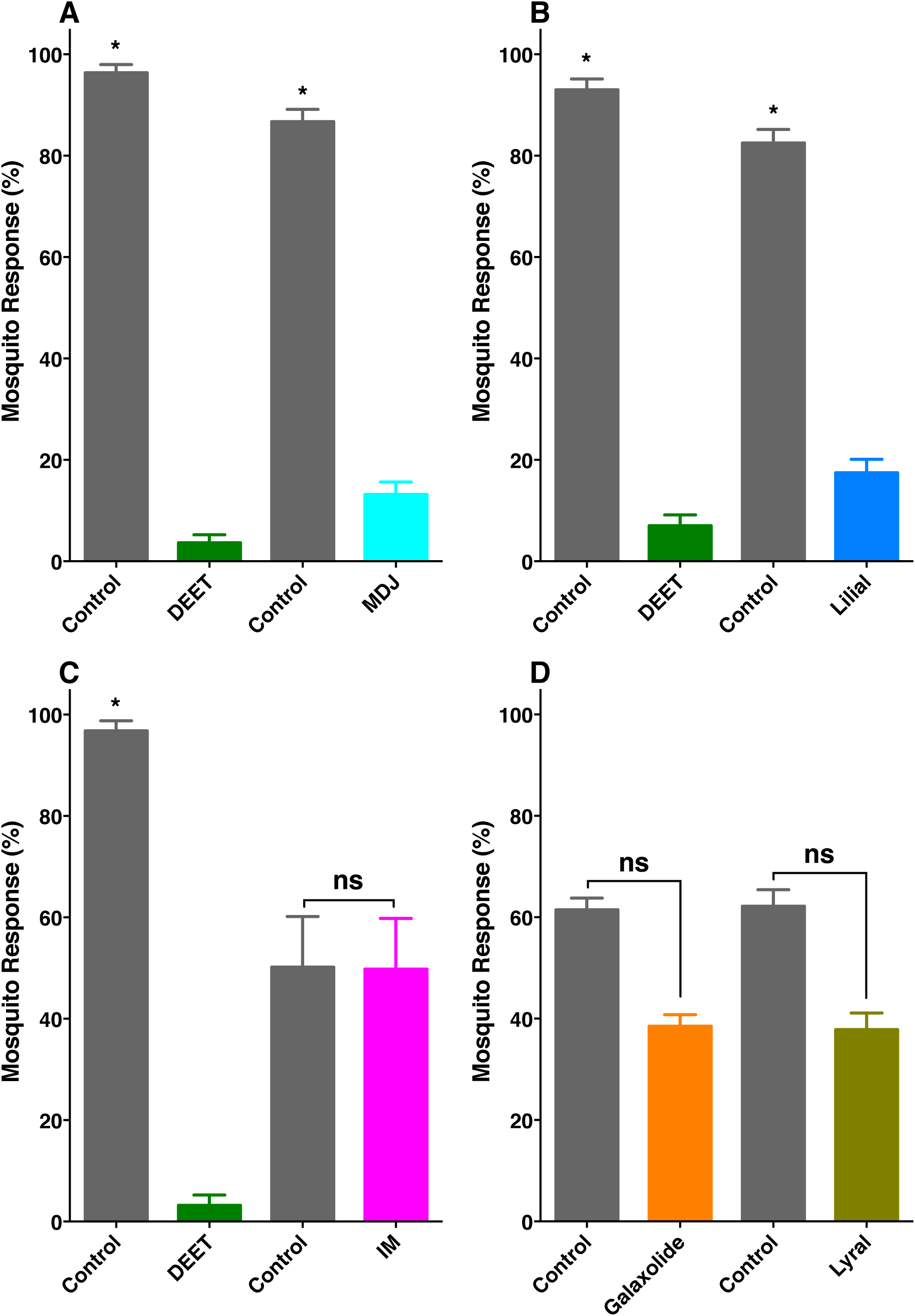
Behavioral responses of blood-seeking female *Culex* mosquitoes to the major constituents of Bombshell in a surface landing and feeding assay. (A) Methyl dihydrojasmonate (MDJ), (B) lilial, (C) isopropyl myristate (IM), (D) galaxolide and lyral – the latter was found in Ivanka Trump eau de parfum. All compounds were tested at 1% dose, and DEET at the same dose was used as a positive control. Data were normalized and expressed as mean ± SEM. Asterisks and “ns” indicate significant (Student *t* test, P < 0.05) and not significant differences, respectively. The number of replicates were (A), DEET, n=12; MDJ, n=11; (B) DEET, n=8; lilial, n=10; (C), DEET and IM, n=4; (D) galaxolide and lyral (n=6).

**Fig. 3.**
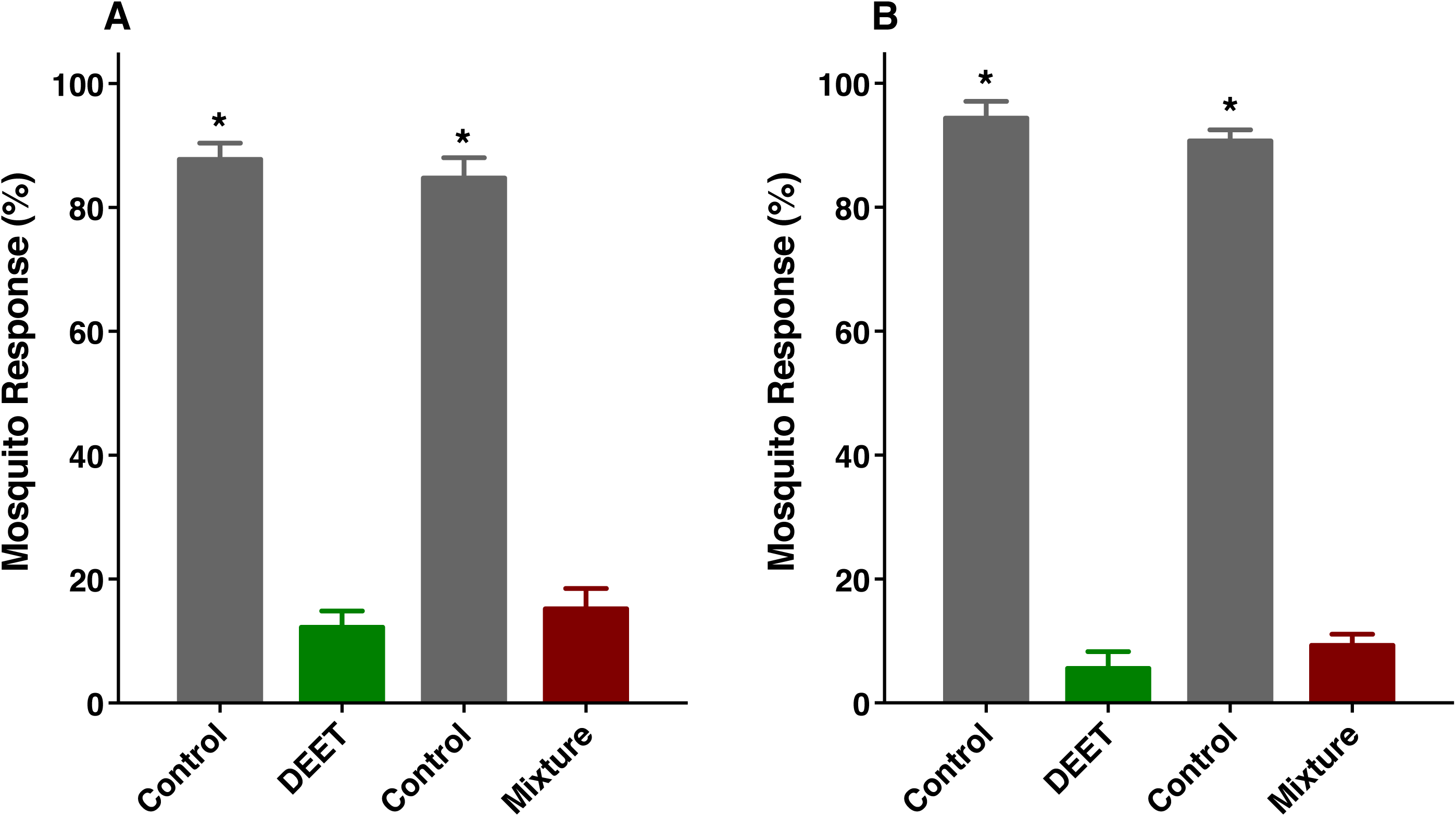
Repellence activity elicited by mixtures of methyl dihydrojasmonate and lilial at (A) 1% and (B) 5% compared with responses to DEET at the same concentration. Data were normalized and expressed as mean ± SEM. Asterisks denote significant differences of transformed data (Student *t* test, P < 0.05). The number of replicates were (A) mixture, n=13; DEET, n=12; (B) mixture and DEET, n=6.

Previously, we identified an odorant receptor from the southern house mosquito, CquiOR136, which is sensitive to mosquito repellents [10]. We expressed CquiOR136 along with its mandatory co-receptor, CquiOrco, in *Xenopus* oocytes and tested their responses to methyl dihydrojasmonate and lilial. Although, both compounds were somewhat strong repellents, they activated CquiOR136 differently. The currents elicited by methyl dihydrojasmonate were significantly higher than those elicited by lilial and even DEET, with all ligands at a 1-mM dose (Fig. 4). We then suggest that the activities of methyl dihydrojasmonate and lilial as spatial repellents were mediated at least in part by CquiOR136.

**Fig. 4.**
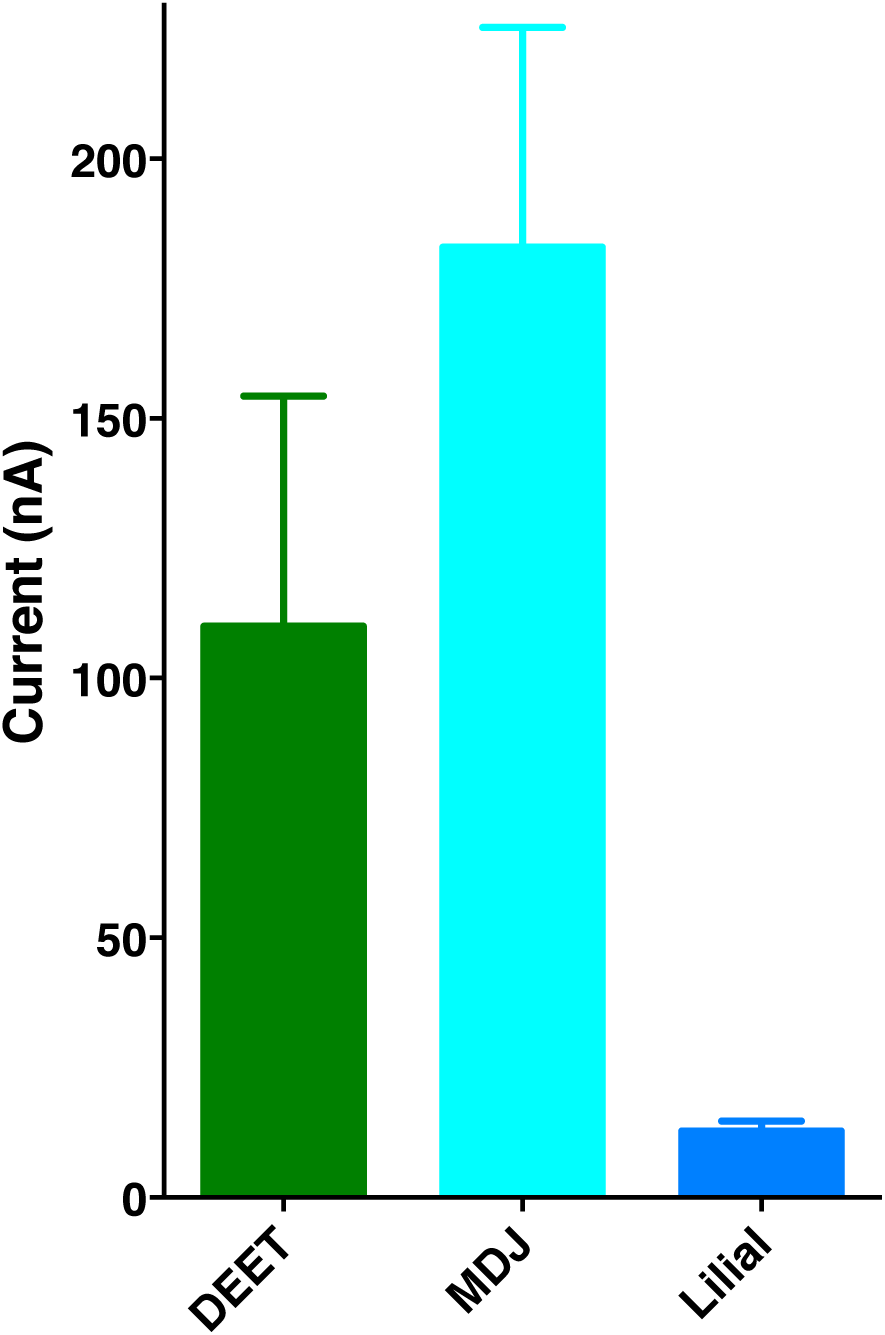
Quantification of current responses elicited by MDJ and lilial on *Xenopus* oocytes expressing CquiOR136/CquiOrco. DEET was applied as a positive control. All compounds were tested at the same dose (1 mM). The data are expressed as mean ± SEM.

Chemical analysis of another perfume, Ivanka Trump eau de parfum, had a similar profile, particularly with regard to the major constituents, except for isopropyl myristate that appeared at much lower levels in the latter perfume (Fig. 1). They also differed in other minor constituents that appeared in Ivanka Trump eau de parfum, but not in Bombshell. Of note, a peak at 13.41 min in the former was identified as lyral based on comparison of MS and retention time obtained with authentic lyral. In our surface landing and feeding assay, lyral demonstrated no repellence activity (Fig. 2D).

Whereas Ivanka Trump eau de parfum has a significantly higher content of methyl dihydrojasmonate than Bombshell has, the content of lilial in the latter was higher than in the former (Fig. 5). Interestingly, the major constituent of these eaux de parfums does not appear in their labels. It might be that the disclosure of constituents in their labels is meant to comply with the Seventh Amendment to the European Cosmetic Directive demanding that cosmetics on sale in Europe indicate whether certain compounds are present at any level [8]. Various minor constituents in these perfumes, as well as lilial and lyral, make the list of 26 compounds; however, methyl dihydrojasmonate is not included. Thus, there is no legal requirement to disclose this compound on labels, despite it being the major constituent in these perfumes.

**Fig. 5.**
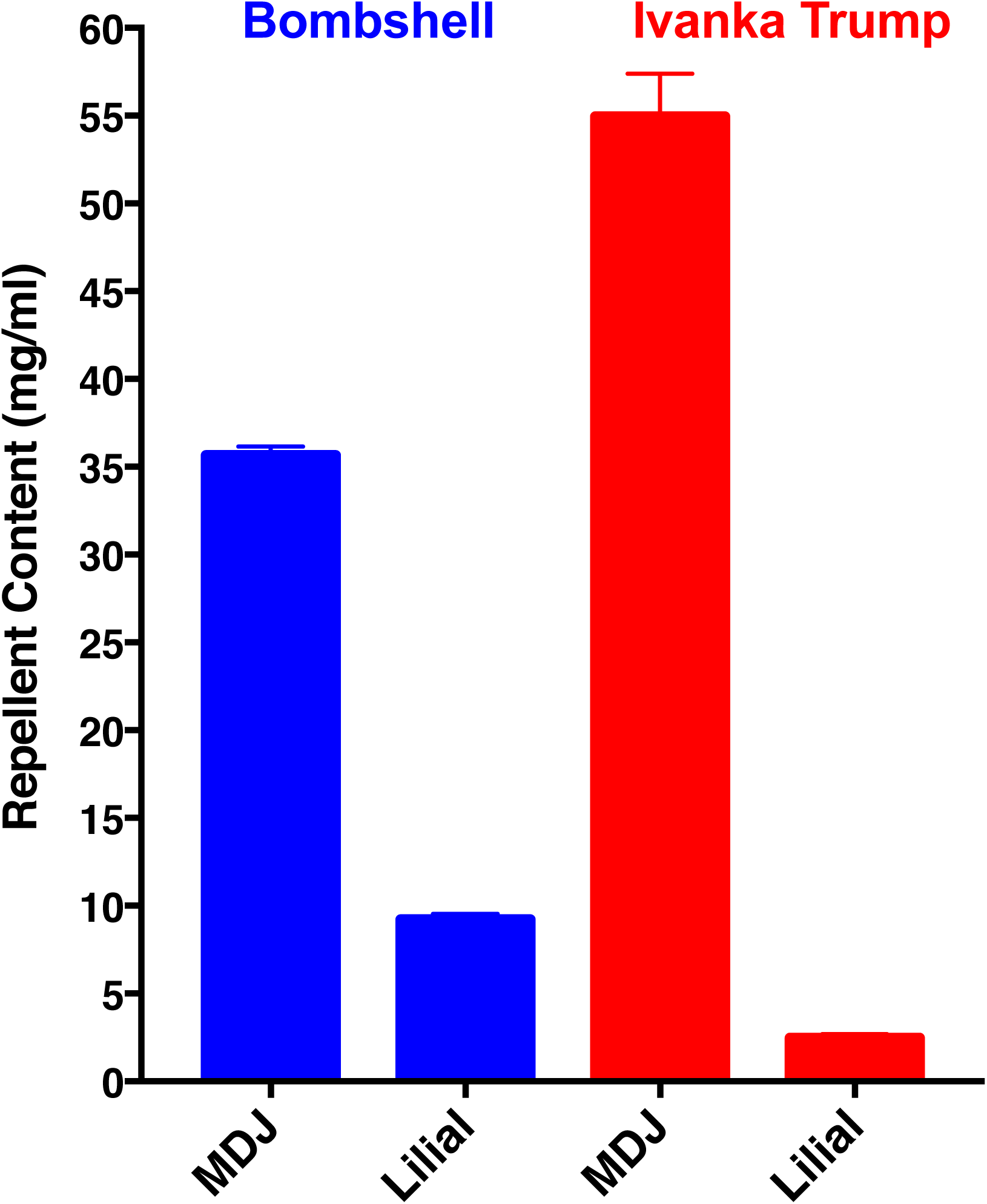
Concentrations of MDJ and lilial in Bombshell (left, blue) and Ivanka Trump (right, red) eaux de parfums. Amounts were estimated by gas chromatography after calibrating the responses of the flame ionization detector with standards. Perfumes were diluted 1,000x for injections (n=3). The data are expressed as mean ± SEM.

A major concern about natural repellents is the complete protection time, i.e., how long they would last as active repellents. Many compounds are misleadingly effective as repellents when tested only at the time the samples are prepared, but not over a reasonable period of time. As opposed to DEET and picaridin, many natural products have an initial spike of activity, because their vapor pressures are very high (low boiling points) thus releasing initially overwhelming doses, but they lose activity over time as the sources are rapidly depleted. In short, even when testing repellents at the same nominal doses, one must keep in mind that the more volatile compounds will have a higher vapor dose initially, whereas the less volatile compounds have lower vapor doses, but they will last longer. DEET has an optimal boiling point for a repellent (545°F = 285°C; PubChem), which allows a steady vapor concentration at the skin surface for a long period of time. Over time, DEET loses activity due to skin penetration and wash off, but loss due to evaporation is minimal [13]. Because perfumes are notorious for depleting over a short duration, despite the new technologies and the availability of fixatives, we asked whether these two eaux de parfums would retain activity for a reasonable period of time. Surprisingly, both Bombshell and Ivanka Trump eaux de parfums retained activity for as long as 6 h (Fig. 6).

**Fig. 6.**
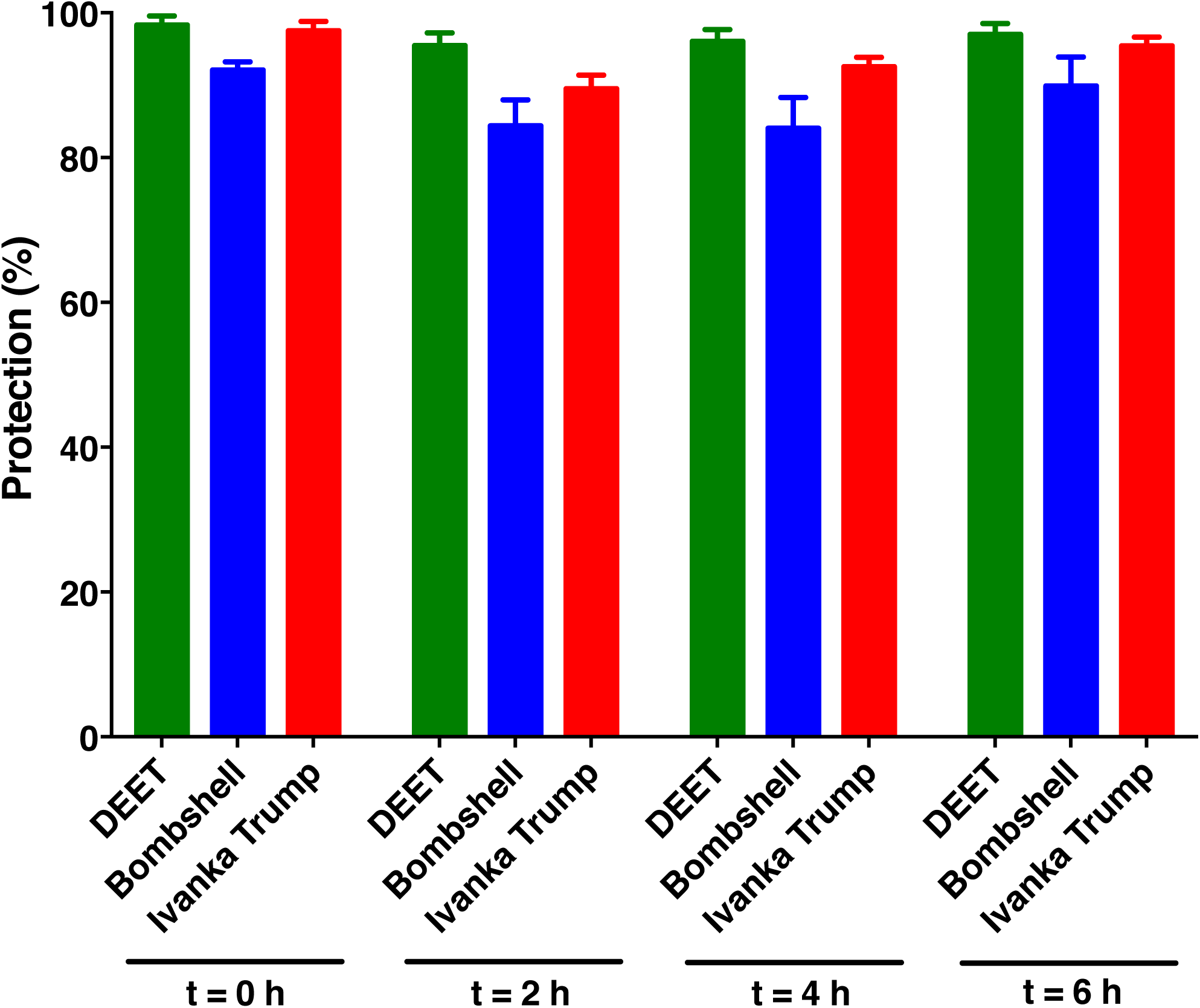
Repellence activity elicited by Bombshell and Ivanka Trump eaux de parfums over a period of six hours. DEET at 5% in our experimental setup, which is equivalent to a commercial formulation of 30% [13], was used as positive control. The perfumes were undiluted in these tests. Data are expressed in protection (%), representing the mean ± SEM of 6 replicates each.

It is worth mentioning that our assays did not measure loses (e.g., wash off, skin penetration) other than loss due to evaporation (from a filter paper; see Material and Methods). Additionally, our tests were conducted with aliquots of 200 μl of each eau de parfum to be consistent with the volume of repellents applied in our repellent assays [13]. Of note, DEET 5% in our experimental setup is nearly equivalent to a commercial formulation with 30% DEET [13]. In our behavioural measurements with 5% DEET, 10 mg of this repellent was used per test. Considering the amounts of methyl dihydrojasmonate in Ivanka Trump eau de parfum, i.e., peak 1, 42.65±1.95 mg/ml and peak 2, 12.26±0.58 mg/ml, we loaded in these comparative assays ≈11 mg of methyl dihydrojasmonate and ≈0.5 mg of lilial (2.46±0.23 mg/ml). Likewise, from Bombshell (peak 1, 30.29±0.49 mg/ml and peak 2, 5.35±0.07 mg/ml), we applied ≈7 mg of methyl dihydrojasmonate and ≈1.8 mg of lilial (9.21±0.32 mg/ml). In short, as far as the amounts of repellents were concerned, these compounds performed nearly equally. It is very important, however, to point out that these comparisons may be misleading as no one applies perfume at levels comparable to repellent applications. Here, 200-μl solutions were applied to approximately 20 cm^2^ [13], but a standard application of DEET is 1 ml of a 20% solution applied to 600 cm^2^ [14]. Thus, in practical applications on the skin, DEET is applied at approximately 0.34 mg/cm^2^. Since Ivanka Trump eau de parfum dispensed 50±2.6 mg of perfume/spray (n=3) and covered an area of the forearm of 24.7±1.5 cm^2^/spray (n=3) [similar results obtained with Bombshell were 55±1 mg/spray and 28.3±0.9 cm^2^/spray] and considering the estimated densities of these perfumes (Ivanka Trump, 0.858±0.18 g/ml; Bombshell, 0.861±0.003 g/ml), the actual amounts of total active repellents in these cosmetics applied to the skin were estimated to be 0.13 mg/cm^2^ (Ivanka Trump) and 0.1 mg/cm^2^ (Bombshell). In other words, even excessive users are unlikely to apply perfume at doses comparable to that of repellents. They typically apply three times lower doses of these perfume-derived mosquito repellents than DEET (from repellent formulations). Although these perfume applications may suffice to fend off nuisance mosquitoes, it might not be a wise prophylactic tool for preventing bites of mosquitoes in areas with arboviruses or other mosquito-borne diseases.

Lastly, we performed a blind test to determine which, if any, of these eaux de parfums would smell good to young people. A blind test was conducted with 18- to 23-year-old students on the UC-Davis campus. They were presented with spray bottles labelled with code names, i.e., Isoleucine/Threonine (IT for Ivanka Trump) and Serine/Histidine (SH, for Bombshell) and asked if they prefer one of them; one column was provided for other remarks. There were no responders that disliked both products; a woman student indicated that “both smell like hand sanitizers,” but she preferred IT. In general, the majority of the responders preferred SH, with more pronounced difference amongst women than men (Fig. 7). Of the 104 students who responded, 62 did not make any remarks, just selected one of the two choices. Some noteworthy remarks were “smell like angel” (IT), “more refreshing” (SH), “love it!” (IT), “more pleasant to smell” (SH), “less harsh” (IT), “sweet” and “sweeter smell” (both SH), “too sweet” (IT), “smell[s] less strong than IT” (SH), “SH smell[s] like hair spray” (IT), “SH smells like flowers, IT smells like vodka” (SH), “smell[s] fruity” (IT), “smells sexy”, “smells like a perfume I already have” (both SH), “amazing!!!” (IT).

**Fig. 7.**
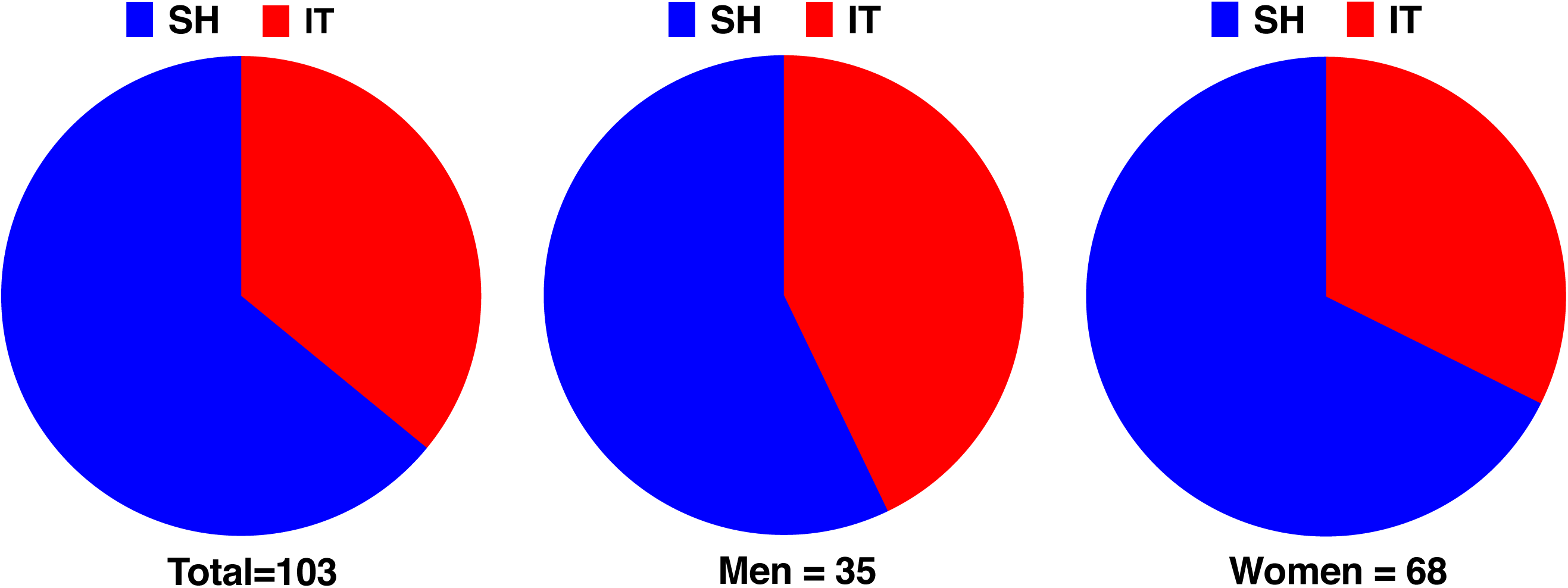
Pie charts summarizing blind preference tests comparing the two eaux de parfums by 18- to 23-year-old students. Bombshell and Ivanka Trump eaux de parfums were provided in spray bottles labelled with code names, i.e., Serine-Histidine (SH) for Bombshell and Isoleucine-Threonine (IT) for Ivanka Trump brand.

## Conclusions

We have identified the active ingredients that make two eaux de parfums, Bombshell and Ivanka Trump, repel blood-seeking *Culex* mosquitoes, i.e., methyl dihydrojasmonate and lilial. Albeit not recommended by us, the “off-label” use of these eaux de parfums as mosquito repellents might help fend off nuisance mosquitoes. However, they might not serve as prophylactic tools against infected mosquitoes for which higher doses of repellents have been recommended [13].

## Acknowledgments

The authors thank Dr. Anthon Cornel (University of California, Department of Entomology & Nematology) for providing mosquitoes that allowed us to duplicate his colony at the Davis campus, students who volunteered for the “smell test,” and Dr. Mona Monfared for allowing us to invite her students to participate in the blind test. F.Z. was supported by a student fellowship from the China Scholarship Council and Huazhong Agricultural University.

## Supporting Information

**S1 File. Dataset for figures.** This file contains the original data that generated figures 2-7.

## References

1. Norris EJ, Coats JR. Current and Future Repellent Technologies: The Potential of Spatial Repellents and Their Place in Mosquito-Borne Disease Control. Int J Environ Res Public Health. 2017;14(2). doi: 10.3390/ijerph14020124. PubMed PMID: 28146066; PubMed Central PMCID: PMCPMC5334678.

2. Moore SJ. Plant-Based Insect Repellents. Insect Repellents Handbook, 2nd Edition. 2015:179-211. PubMed PMID: WOS:000355003300010.

3. Gross AD, Coats JR. Can Green Chemistry Provide Effective Repellents? Insect Repellents Handbook, 2nd Edition. 2015:75-90. PubMed PMID: WOS:000355003300006.

4. EPA. DEET 2016 [cited 2017 July 21]. Available from: https://www.epa.gov/insect-repellents/deet.

5. Strickman D. Topical Repellent Active Ingredients in Common Use. Insect Repellents Handbook, 2nd Edition. 2015:231-7. PubMed PMID: WOS:000355003300012.

6. Patel RV, Shaeer KM, Patel P, Garmaza A, Wiangkham K, Franks RB, et al. EPA-registered repellents for mosquitoes transmitting emerging viral disease. Pharmacotherapy. 2016;36(12):1272-80. doi: 10.1002/phar.1854. PubMed PMID: 27779781.

7. Gu X, Wang T, Collins DM, Kasichayanula S, Burczynski FJ. In vitro evaluation of concurrent use of commercially available insect repellent and sunscreen preparations. Br J Dermatol. 2005;152(6):1263-7. doi: 10.1111/j.1365-2133.2005.06691.x. PubMed PMID: 15948991.

8. Sell CS. Chemistry and the Sense of Smell. Chemistry and the Sense of Smell. 2014:1-459. doi: 10.1002/9781118522981. PubMed PMID: WOS:000351669100013.

9. Rodriguez SD, Drake LL, Price DP, Hammond JI, Hansen IA. The Efficacy of Some Commercially Available Insect Repellents for *Aedes aegypti* (Diptera: Culicidae) and *Aedes albopictus* (Diptera: Culicidae). J Insect Sci. 2015;15:140. doi: 10.1093/jisesa/iev125. PubMed PMID: 26443777; PubMed Central PMCID: PMCPMC4667684.

10. Xu P, Choo YM, De La Rosa A, Leal WS. Mosquito odorant receptor for DEET and methyl jasmonate. Proceedings of the National Academy of Sciences of the United States of America. 2014;111(46):16592-7. doi: 10.1073/pnas.1417244111. PubMed PMID: 25349401; PubMed Central PMCID: PMC4246313.

11. Chapuis C. The Jubilee of Methyl Jasmonate and Hedione (R). Helv Chim Acta. 2012;95(9):1479-511. doi: 10.1002/hlca.201200070. PubMed PMID: WOS:000308718200002.

12. NIST. NIST Chemistry WebBook 2017 [cited 2017 July 20]. Available from: http://webbook.nist.gov/cgi/cbook.cgi?ID=C80546&Mask=80#IR-Spec.

13. Leal WS, Barbosa RM, Zeng F, Faierstein GB, Tan K, Paiva MH, et al. Does Zika virus infection affect mosquito response to repellents? Sci Rep. 2017;7:42826. doi: 10.1038/srep42826. PubMed PMID: 28205633; PubMed Central PMCID: PMCPMC5311973.

14. WHO. Guidelines for efficacy testing of mosquito repellents for human skin 2009 [cited 2016 November 14]. Available from: http://apps.who.int/iris/bitstream/10665/70072/1/WHO_HTM_NTD_WHOPES_2009.4_eng.pdf.

